# Transgene Expression Kinetics and Replication Potential of Recombinant Adenovirus Serotype 4 in a Mouse Model and its Use as a Herpes Simplex Virus Vaccine

**DOI:** 10.64898/2026.05.15.725395

**Authors:** Alexander C. Vostal, Dawid Maciorowski, James M. Readler, Isabella S. Pytel, Andy Patamawenu, Ceanna Cooney, Peyton M. Roeder, Rachel Roenicke, Freya van’t Veer, Tae Kim, Ellison Ober, Yuyan Yi, Jingwen Gu, Mitra Harrison, Breanna Kim, Guangping Liu, Kennichi Dowdell, Anna Hostal, Kening Wang, Mark Connors, Jeffrey I. Cohen

**Author notes:** Corresponding Author: Alexander C. Vostal, MD. Medical Virology Section, Laboratory of Infectious Diseases, National Institute of Allergy and Infectious Diseases, 50 South Dr, Bethesda, MD 20894, USA. **Alternate Corresponding** Author: Jeffrey I. Cohen, MD.

## Abstract

Human adenovirus serotype 4 (Ad4) is used as a replication-competent oral vaccine that safely and effectively prevents Ad4 respiratory illness in US military personnel. Recombinant Ad4 vaccine candidates elicit mucosal and systemic immune responses against respiratory viruses in hamsters, nonhuman primates, and humans. Although evaluation of Ad4 vaccine candidates in mice would be extremely useful given the large number of immunologic tools available, this has been limited by concerns about a lack of viral replication in these animals. Here we generated recombinant Ad4 vectors that express either luciferase (Ad4-Luc) or herpes simplex virus type 2 (HSV-2) glycoprotein D (Ad4-gD2) to identify transgene expression kinetics, the presence of Ad4 vector replication, and HSV-2 immune responses and protection against HSV-2 infection. Local luciferase activity was observed from 7 hours to 20 days after intranasal inoculation of BALB/c and humanized mice. Subsequent inoculations with Ad4-Luc showed reduced luciferase expression in BALB/c mice, but robust expression in humanized mice, suggesting an immune response to the vector in wild-type mice. Ad4 DNA, but not luciferase activity, was reduced in the lungs of BALB/c mice treated with cidofovir before inoculation with Ad4, implying that Ad4 replicated, albeit at a low level, in the lungs. Intranasal vaccination of mice with Ad4-gD2 resulted in HSV-2 neutralizing antibody in the serum, and after HSV-2 intravaginal challenge reduced disease scores, increased survival, and reduced shedding. Overall, the BALB/c mouse model is semi-permissive to Ad4 mucosal infection, but transgene expression is sufficient for the study of Ad4-based vaccine candidates.

**Importance:** Mucosal surfaces serve as the primary site of infection and shedding for many viral pathogens. Immune responses at mucosal sites provide protection, but few mucosal vaccines are licensed. The oral replication-competent adenovirus serotype 4 (Ad4) vaccine is used to prevent respiratory illness in military recruits, has an extraordinary record of safety and efficacy and has been tested as a recombinant platform for other viruses. Further development of this vaccine platform has been partially hindered by the perceived inability to evaluate vaccine candidates in mice. Here we characterize recombinant Ad4 transgene expression kinetics and viral replication after inoculation at various sites and show protection against herpes simplex virus type 2 (HSV-2) genital disease in mice after intranasal vaccination. We show that Ad4 can induce protective efficacy, even in a semi-permissive mouse model, suggesting this is a promising vector for HSV-2 and potentially other viral pathogens.

## INTRODUCTION

Human adenoviruses (HAdV) are double-stranded DNA viruses that cause respiratory infections with high worldwide seroprevalence ^1–3^. Frequent adenovirus serotype 4 (Ad4) outbreaks in US military recruits were documented in the 1950’s and 1960’s, leading to the development of a live, enteric-coated bivalent vaccine ^2,4–7^. Clinical trials demonstrated that selective infection of the gastrointestinal tract with a wild-type Ad4 or Ad7 isolate in an enteric coated pill almost entirely prevented febrile respiratory illness and hospitalization due to the virus, resulting in FDA approval of Ad4/Ad7 bivalent vaccine in 1980 for all military personnel^8,9^. The effectiveness of this vaccine led to a vaccination program for military recruits from 1971 to 1999. Vaccination was paused for the next 10 years when the manufacturer stopped production and then resumed in 2011 and continues to the present with over 10 million doses administered and an extraordinary safety record^8,10^. The immunogenicity and protection elicited by this replication-competent adenovirus vaccine suggests that it might also be effective as a vaccine vector to express other viral immunogens.

Many other recombinant adenoviral vectors have been used to elicit immune responses to other pathogens in preclinical studies and clinical trials, but most have relied on replication-incompetent vectors which may limit their immunogenicity and effectiveness ^11^. Matsuda et. al. evaluated a replication-competent recombinant Ad4 vector expressing influenza H5 haemagglutinin and demonstrated its ability to safely induce long lasting systemic and mucosal immune responses in humans^12,13^. The ability of the Ad4 vector to replicate in human volunteers for 2 to 4 weeks has been proposed as an important mechanism in the development of robust long-lasting immunity^12,13^.

HAdV replicates poorly in most mouse cell lines, which has limited the utility of murine models to evaluate replication-competent HAdV vectors ^14,15^. Mice do not express CD46 which is an essential entry receptor for group B and D HAdVs^16–19^. Group E HAdVs, which include Ad4, can enter most mouse cell lines and effectively express early genes but are often unable to replicate in these cells^15,20^. Viral replication may also be limited in mice due to the suppression of adenovirus E1 genes and degradation of proteins necessary for adenovirus replication in mouse cells^20–26^. Rodriguez et. al. generated a humanized mouse model of HAdV infection by transplanting NSG-A2 mice with CD34+ human hematopoietic stem cells and challenged the animals intravenously with HAdV-C2 ^27^. While over a third of the mice developed acute infection, it was unclear if HAdV-C2 was replicating in the human or mouse cells. Furthermore, Chen et.al. recently demonstrated that BALB/c mice infected intravenously with replication-competent Ad5 develop organ pathology and mortality which may be helpful as a model to study adenovirus-based therapeutics^28^. Although the extent of adenovirus replication in mice remains unconfirmed, two groups have demonstrated transgene expression and immunogenicity of a recombinant replication-competent Ad4 vaccine vector in murine models ^29 30^. However, it was unknown if the immunogenicity was due to transgene expression in the presence or absence of adenovirus replication. In view of the concerns about limited replication of Ad4 in mice, hamsters have been used more often as a model for preclinical development of vectored Ad4 vaccines^31 32^. In this model, shedding of Ad4 viral DNA has been observed for at least 5 days after intranasal inoculation^31^, which is less than the 2 to 4 weeks of shedding observed in humans^12^, but sufficient to induce potent immune responses. However, immunological reagents are limited for hamsters in comparison with mice.

To study transgene expression kinetics with a recombinant Ad4 vector in mice, we generated a replication-competent Ad4 vector that expresses luciferase (Ad4-Luc) and followed luciferase expression kinetics in mice that were inoculated with Ad4-Luc at different sites. We found consistent luciferase expression at the site of inoculation and in distant organs in mice infected with Ad4-Luc. Ad4 DNA was reduced in lung tissue of mice treated with the antiviral cidofovir. To establish the feasibility of Ad4 as a vaccine vector in mice, we produced an Ad4 expressing herpes simplex virus type 2 (HSV-2) glycoprotein D (gD2). We utilized different routes of mucosal inoculation to study the immunogenicity of Ad4-gD2 in mice and demonstrated its ability to protect mice from lethal HSV-2 challenge.

## METHODS

### Production of Recombinant Replication-Competent Ad4 vectors

Adenoviruses expressing transgenic proteins, enhanced green fluorescent protein (Ad4-EGFP; Synthetic IDT DNA Technologies) and Luciferase (Ad4-Luc; GenBank Accession #AB762768.1), were constructed using a previously reported modified Ad4 genome sequence by Alexander et. al.^30^. A pUC19 shuttle vector with a cassette in the multiple cloning site containing EGFP or Luc with a cytomegalovirus (CMV) 433 promotor and Kozak sequence (GCCACC) at the 5’ end and a bovine growth hormone polyadenylation signal (BGH polyA signal) at the 3’ end **[Figure 1a]** flanked by two regions of homology to the E3 region (684 bp; nucleotide positions 28880-29563 and 592 bp; 32207-32798) of the Ad4 genome was constructed. A plasmid containing the entire modified Ad4 genome (including a kanamycin expression cassette for recombination (2,570 bp which crosses the end of the circularized genome; nucleotide position 35751-2675), or “Ad4 backbone,” was cut at restriction enzyme sites, SpeI (NEB, cat. R3133S), (nucleotide positions 30855 and 30865 of the modified Ad4 genome within the E3 region). The previously described shuttle vector was cut with HindIII-HF (NEB, cat. R3104S) and SacII (NEB, cat. R0157S), removing the ampicillin cassette. For homologous recombination, 50-100 ng of the purified linearized Ad4 backbone was combined with 100-200 ng of purified linearized pUC19-EGFP or pUC19-Luc in Eppendorf tubes on ice. 40 ul of electro-competent E. coli BJ5183 strain (RecBC) was added to each Eppendorf tube and then transferred to 0.2 cm cuvettes. The bacteria and DNA mixture were electroporated (Bio-Rad Micropulser) at 2.5 kv. SOC media was added to the cuvette, and cells were transferred to a Falcon 2059 culture tube to incubate at 37°C, and shaken at 225 rpm for 1 hour. After 1 hour, the cells were pelleted (3000 rpm, 10 min), plated onto kanamycin (50 μg/ml) agar plates, and grown overnight at 37°C. Bacterial colonies were picked and screened by PCR with the Ad4 CMV promoter forward primer (5’- TCATCAAGACCCTATGCGG-3’) and Ad4 Fiber (L4) reverse primer (5’ GAATCCATCTGAAGAGACGAAGGG-3’), using a Q5 High-Fidelity PCR enzyme kit (NEB, cat. M0491S), following the manufacturer’s protocol. PCR positive colonies were grown overnight and DNA was digested with BamHI-HF to identify Ad4 containing either EGFP (pRAd4-EGFP) or Luc (pRAd4-Luc). PCR positive BJ5183 miniprep DNAs were retransformed again in E.coli strain NEB10-recBA, using the manufacturer’s protocol (New England Biolabs, cat. C3019H). Miniprep DNAs were prepared and DNA inserts were verified by sequencing and the plasmids were grown as Maxiprep DNA using a Qiagen Maxi-Endo-Free kit (cat. 12362).

**Fig. 1.**
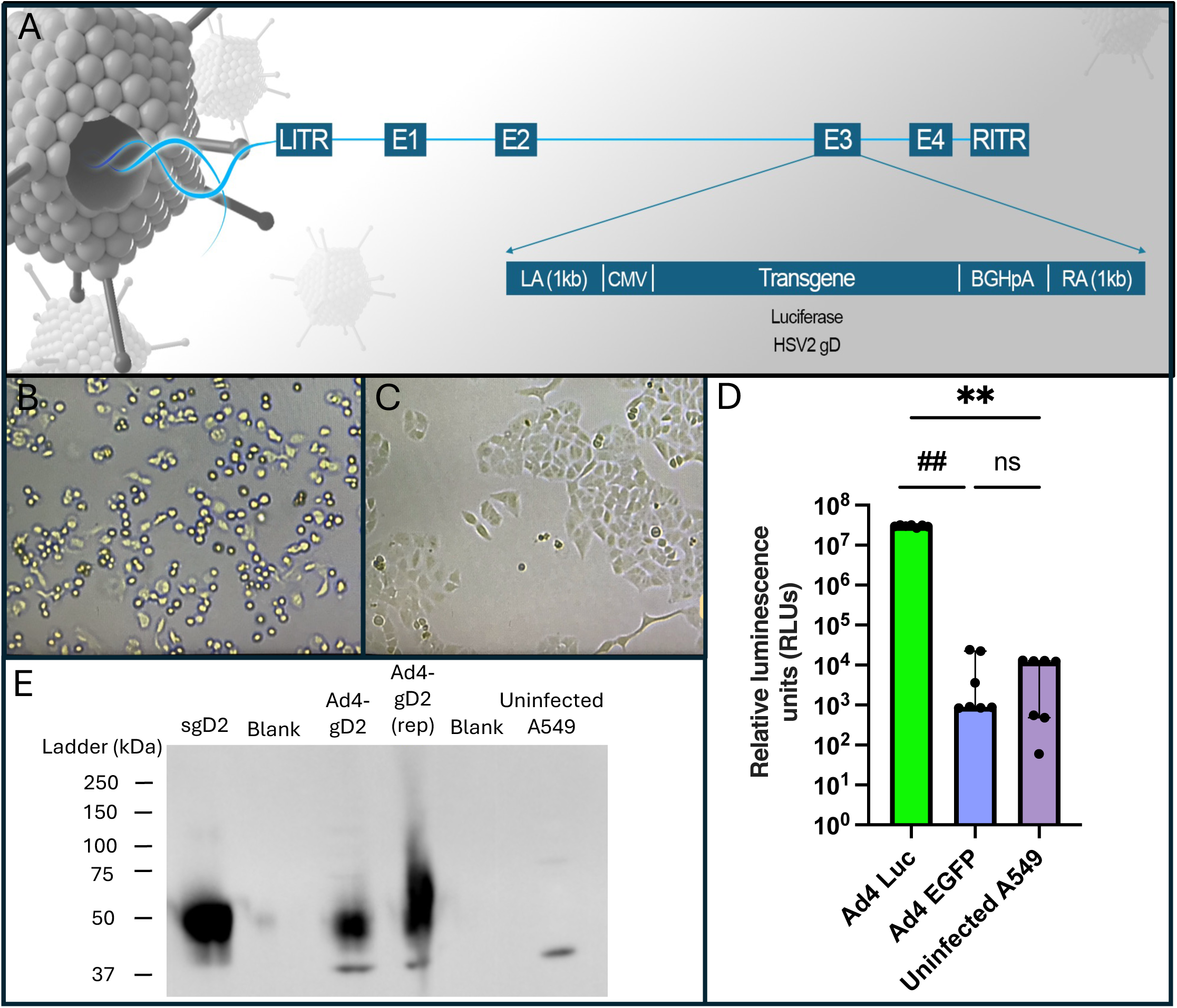
Construction and characterization of Ad4 expressing luciferase or HSV-2 gD. (A) The E3 region of the adenovirus serotype 4 (Ad4) genome is replaced by left and right arms (LA, RA) to allow recombination with a cassette with LA, a CMV promoter, a luciferase or HSV-2 gD gene, a bovine growth hormone (BGH) poly A site, and an RA. The recombinant Ad4 genome is approximately 37 kb in length and retains genes in the E1 region responsible for viral replication. The genome map abbreviations are as follows; left inverted terminal repeat (LITR), E1-early transcriptional gene region 1 (consisting of E1A and E1B), E2-early transcriptional gene region 2 (consisting of E2A and E2B), E3-early transcriptional gene region 3, E4-early transcriptional gene region 4, and right inverted terminal repeat (RITR). (B) In vitro Ad4-Luc infection of A549 cells showing cytopathic effects (MOI 0.1, 3 days after infection, 5x magnification). (C) Uninfected A549 cells for comparison (5x magnification). (D) Relative luminescence of A549 cells infected with Ad4-Luc, Ad4-EGFP, or uninfected. Bars show median values with interquartile range (IQR), with individual data points. ## p=0.009 (Ad4-Luc versus Ad4-EGFP); ** p=0.002 (Ad4-Luc versus uninfected); ns: not significant. (E) Western blot of purified soluble HSV-2 gD protein (Sol gD2), cell lysates of A549 cells infected with Ad4-gD2 (Ad4-gD2), a replicate lysate (Ad4-gD2 rep), and uninfected A549 cells after staining with anti-HSV-2 gD antibody.

The previously described SpeI site was utilized to linearize pRAd4-EGFP or pRAd4-Luc and this was transfected using PEIpro® (Polyplus Cat# 101000033) into the A549 human carcinoma lung cell line (ATCC cat. CCL-185). A549 cells were incubated (37°C and 5% CO_2_) for 14 days, until cytopathic effects were evident **[Figure 1b]**, to produce a primary stock of recombinant Ad4 (Ad4-EGFP or Ad4-Luc). This primary viral stock was used to inoculate mice for Ad4-Luc transgene experiments.

Recombinant Ad4 expressing the full-length of HSV-2 glycoprotein D (gD2) (amino acids 1 to 393) from HSV-2 strain 333 was constructed using Gibson Assembly cloning. Briefly, HSV-2 gD was amplified by PCR using primers (Forward 5’-GAAATCGGAAAGCGGACGCGGAACTAGTGCCCGGGCGCCACCATGGGGCGTTTGACCTCC-3’ and Reverse 5’-CAAGCGCTCACGGGATACTCGACTAGTGCCCGGGCCTAGTAAAACAATGGCTGG-3’) from a CMV expression plasmid containing the HSV-2 gD nucleic acid sequence. The original Ad4 backbone was modified (2^nd^ generation Ad4 backbone) to accommodate Gibson Assembly cloning. Briefly, the nucleic acid sequence for the entire E3 protein, including approximately 100 nucleotides upstream of the 5’ end of the E3 protein, was removed via bacterial homologous recombination. Modified One-step Site (MOS) sequences were added downstream from the previously described 5’ CMV promotor and upstream from the 3’ BGH polyA signal. 120 ng of the amplified gD2 sequence was combined with 100 ng of linearized 2^nd^ generation Ad4 backbone and the mixture was incubated with Gibson Assembly^®^ Master Mix (NEB, cat. E2611L) for 30 min at 50°C, added to chemically-competent NEB10-beta cells on ice, and heat-shocked for 30 seconds at 42°C. The cells were grown in SOC media and plated on kanamycin agar plates. Bacterial colony screening was performed as described above and the sequence of the Ad4 backbone containing gD2 was confirmed with Sanger sequencing. The recombinant Ad4-gD2 genome was linearized with PacI and then transfected and propagated in A549 cells as described above. A primary stock of Ad4-gD2 was further purified using an ion exchange column (Sartobind Q capsule 75 mL bed volume, Sartorius, Bohemia, NY), ultra-filtrated (Midi 20cm 500kD mPES 0.5mm, Repligen, Waltham, MA) ^30^, and used for vaccination of mice.

Purified Ad4 was titrated in A549 cells and infection forming units (IFU) were determined by incubating cells with serial dilutions of virus for 24 to 48 hours, fixing the cells, staining the cells with mouse anti-adenovirus hexon antibody followed by horseradish peroxidase (HRP)-conjugated goat anti-mouse secondary (AdEasy Viral Titer Kit, Cat *#*972500, Agilent), and then counting the number of infected cells at 5x magnification. In vitro expression of luciferase was confirmed by infecting A549 cells with Ad4-Luc at MOI of 0.1, and three days after infection the cells were treated with lysis buffer (Promega 5x cell culture lysis solution, Cat.# E1500), luciferase assay reagent (Cat.# E1500) was added, and luciferase expression of the cell lysate was quantified in a 96-well black plate using a PerkinElmer Victor X2 Multilabel Microplate Reader. In vitro expression of HSV-2 gD was confirmed by collecting cells three days after Ad4-gD infection, freeze-thawing the cells, and adding RIPA lysis and extraction Buffer (Thermo-Fischer). Western blotting was performed on cell lysates, and the blot was incubated with rabbit anti-gD2 antibody (R7)^33^, followed by goat anti-rabbit IgG HRP-linked secondary antibody.

### Adenovirus Inoculations

Mouse studies were performed under a protocol approved by the Animal Care and Use Committee of the National Institute of Allergy and Infectious Diseases (NIAID). For Ad4-Luc mouse infections, male and female BALB/c mice (Charles River Laboratories or from an NIAID breeding colony) were obtained between 4 to 13 weeks old. CD34+ NCG or NOG humanized mice (HuCD34+ NCG, Charles River Laboratories and HuCD34+ NOG, Taconic) were obtained at an average of 24 weeks (range 18 to 26 weeks) after human CD34+ cord blood cell infusion. Mice were inoculated with Ad4-Luc intranasally (IN, 20 ul in each nostril), intramuscularly (IM, 50 ul), via oral gastric gavage (OG, 200 ul) or intravaginally (Ivag, 20 ul). Isoflurane was used for sedation prior to IN inoculation. Ad4 given via OG was diluted in 2% bicarbonate solution with or without addition of protease inhibitor aprotinin (final concentration 1 mg/ml, EMD Millipore #616370) or in phosphate buffered saline (PBS) based on prior studies^34,35^. Cidofovir was purchased as a powder (Selleckchem.com, Cat: S1516) or in solution (75 mg/ml Avet Pharmaceuticals Cat: 021631) diluted with saline and administered to mice intraperitoneally at 100 mg/kg.

For Ad4-gD2 mouse infections, 7-week-old female BALB/c mice (Envigo-RMS) received medroxyprogesterone acetate (3 mg) subcutaneously 8 days before inoculation with purified Ad4-gD2 (1.0 × 10^13^ IFU/mL). Medroxyprogesterone was used to thin the vaginal mucosa, to increase susceptibility to Ad4-gD2 infection, and to synchronize the estrous cycle of the mice before Ivag inoculation. Medroxyprogesterone acetate was used for all routes of Ad4-gD2 inoculation to reduce any differences due to the drug. Three routes (n=5 per group) of inoculation were compared: IN, Ivag, and OG (in which Ad4-gD2 was mixed with aprotinin and a 2% bicarbonate solution). Control groups of mice received IM saline and, in the case of Ad4-gD2 experiments, soluble gD2 (5 ug) mixed with Sigma Adjuvant System (Cat# S6322, Sigma-Aldrich) was added as another control. Two doses of each vaccine candidate were administered per group, separated by approximately 3 weeks.

### In vivo and ex vivo imaging of luciferase expression

In vivo imaging of live mice was performed using an IVIS Spectrum In Vivo Imaging System (Caliper Life Sciences) as previously described^31^. Mice were inoculated intranasally with 40 ul (20 ul each nostril) of 1.4 ×10^8^ IFU/mL of Ad4-Luc. Luminescence was measured in anesthetized mice (9 to 34 weeks old at first inoculation) after intraperitoneal injection of D-Luciferin (final dose: 0.15 mg, IVISbrite D-Luciferin Potassium Salt Bioluminescent Substrate, Revvity Health Sciences, Cat#122799) in saline approximately 10 min prior to IVIS imaging. Luminescence was quantified as the average radiance (photons/sec/cm^2^/steradian) using Living Image Software (Revvity Health Sciences, version 4.3.1), and unadjusted values were plotted. A median average radiance in uninfected mice was determined after intraperitoneal injection of D-luciferin for both BALB/c (1,250 average radiance) and HuCD34 (1,865 average radiance) mice **(Supplemental Fig. 1A,B)**. A threshold value, 2-fold higher than the median average radiance value in uninfected BALB/c (2,500 average radiance) and HuCD34 (3,730 average radiance), was used to establish the limit of detection. Mice were given a subcutaneous dose of medroxyprogesterone acetate 8 days before Ad4-gD2 (regardless of the route) or Ad4-Luc (Ivag route only).

For ex vivo imaging of the individual organs, animals were given an intraperitoneal injection of 0.15 mg of D-luciferin and immediately euthanized via CO_2_ inhalation followed by cervical dislocation. Organs, harvested within 20 minutes of euthanasia, were transferred to a clear multi-well dish containing a solution of 300 ug/ml of D-luciferin. The organs were then imaged for luminescence (both average radiance and maximum luminescence over 5 mins) in the IVIS Spectrum In Vivo Imaging System approximately 20 mins after the animals had been euthanized. A threshold value for the limit of detection was determined based on luminescence values per individual organ that was 2-fold higher than the compiled median luminescence (818 average radiance) of all ex vivo organs from uninfected mice **(Supplemental Fig. 1C)**. Similarly, a threshold value for the limit of detection that was 2-fold higher than the median average radiance from uninfected ex vivo lungs (434 average radiance) was also used for IVIS evaluation of isolated ex vivo lung tissue **(Supplemental Fig. 1D)**.

### Quantitation of adenovirus DNA from lung tissue

At necropsy, the left lung was frozen at -80_°_C until used for Ad4-Luc DNA extraction. After thawing, lung samples (ranging from 7 mg to 12 mg) were placed into 2 mL Sarstedt tubes containing 180 µL ATL buffer (Qiagen) and 20 µL proteinase K (lysis buffer). Samples were incubated overnight in a thermomixer at 56_°_C, shaking at 650 rpm. DNA was extracted from 20 ul of lysis buffer (1/10 of total lysis buffer volume) using a DNeasy blood and tissue kit (Qiagen Cat. 69504) and eluted with 20-80 µL molecular biology grade water. DNA was diluted to 10 ng/ul with molecular biology grade water. PCR primers and probes were designed to amplify the adenovirus hexon gene (Forward primer: 5’- GAAACACTGAACTGTCCTACCA-3’, Reverse primer: 5’- TGCGCACATCAGGATCATAG -3’ and Probe: 5’-56-FAM/TGTCCACCG/ZEN CCTGATTCCACATAC/3IABKFQ – 3’). Amplification was carried out in a 20 µL reaction containing 10 µL of Taqman Fast Advanced Master Mix (Applied Biosystems), 2 uL Taqman probe (2.5 µM), 300 nM of each hexon primer, and 40 ng of extracted DNA in 5 uL of molecular biology water. Real-time quantitative PCR was performed using a QuantStudio 6 Flex system (Applied Biosystems). PCR conditions were one cycle of incubation with uracil-DNA glycosylase at 50°C for 2 min followed by 95°C incubation for 20 sec and then 40 cycles of denaturation at 95°C for 1 sec and annealing at 60°C for 20 sec. Analysis was done using QuantStudio 6 Real-Time PCR Software v1.7.2 (Thermo Fisher Scientific). A standard curve was generated by plotting duplicates of PCR amplification of 1:10 dilutions of Ad4-Luc DNA. Each PCR reaction was performed in duplicate and DNA copy number was calculated per mg of lung tissue for each animal.

### Adenovirus Serotype 4 Serum Neutralization Titration

Mouse serum was incubated at 56_°_C for 1 hour to inactivate complement. 5-fold serial dilutions of serum were incubated with Ad4-Luc (MOI=0.01) for 30 min and then added to A549 cells (1.5 × 10^4^ cells/well) in a black 96-well plate overnight at 37_°_C and 5% CO_2_. Cells were then lysed in 1x Cell Culture Lysis Buffer Reagent (Promega, Madison, WI, USA) and luminescence was measured by a PerkinElmer Victor X2 Multiplate reader after addition of 50 ul of luciferin substrate (Luciferase 1000 Assay System, Promega, E4550). Neutralization titrations were calculated on GraphPad software.

### HSV gD ELISA

Serum was obtained from mice 1 week after the second inoculation of Ad4-gD2, soluble gD2 or PBS. Immulon 4 HBX plates (Thermo Fisher Scientific) were coated with 1 µg/mL recombinant soluble gD2 in PBS and incubated overnight at 4°C. Plates were then washed with PBS with 0.05 % Tween and blocked with blocking buffer (2% bovine serum albumin in PBS) for 1 hour at room temperature. Sera were added to the plates in duplicate, serially diluted 2- to 5-fold and then incubated at room temperature for 1 hour. Plates were washed, incubated with anti-mouse HRP-conjugated IgG (Sigma-Aldrich, A9044) for 1 hour, washed, and developed with (3,3’,5,5’-tetramethylbenzidine) TMB substrate (KPL) for 5 min. The reaction was then stopped with 1 M sulfuric acid and absorbance at 450 nm was measured by a spectrophotometer. Endpoint titers were defined as the last dilution to be 2-fold above the average background OD450.

### HSV-2 Neutralization Assay

Vero cells (ATCC) were maintained in DMEM with 5% (vol/vol) fetal bovine serum (FBS), 100 U/mL streptomycin, and 100 u/mL penicillin at 37°C. Sera were heat-inactivated (30 min at 56°C) and then serially diluted into 96 well plates (Corning) followed by adding 70 plaque forming units of HSV-2 R519 (clone of HSV-2 strain 333) and then incubating for 1 hr at room temperature. Then, the virus-serum mixture was added to Vero cells in 6-well plates and incubated at 37°C for 1 hr. The inoculum was removed and DMEM (Gibco) containing 5% (vol/vol) FBS and 0.3% (vol/vol) Gamunex-C (GRIFOLS) was added and then incubated for 2 days at 37°C. Cells were then stained (10% vol/vol formaldehyde, 5% vol/vol acetic acid, 60% vol/vol methanol, and 1% (wt/vol) crystal violet) for 1 hr. After the hour, plates were washed with water and plaques were counted under a microscope. Neutralizing titers were plotted as the serum dilution that inhibited 50% of virus and was calculated with a non-linear regression using GraphPad Prism (Software v.9.3.1).

### HSV-2 Mouse Vaginal Challenge

Mice were injected subcutaneously with 2 mg of medroxyprogesterone acetate 5 days before HSV-2 challenge. A dose of 6.4 × 10^4^ PFU of HSV-2 (strain 333) was administered Ivag on day 0 and vaginal swabs were obtained 1-7 days after challenge and stored at -80°C. Mice were monitored for at least two weeks post challenge and underwent clinical scoring: 0, normal; 1, perineum with mild erythema or some hair loss; 2, perineum with moderate erythema or hair loss or edema; 3, perineum with more than one of the following: erythema, edema, hair loss; 4, perineum with more than two of the following: erythema, edema, wet, hair loss; or mild systemic symptoms like hunched posture, mild hindlimb paresis; 5, perineum disease plus hindlimb paralysis or lethargy; 6, death. Mice were euthanized when they reached clinical end points such as inability to eat or drink, breathing problems or signs of encephalitis (seizures, convulsions, ataxia).

### Data Analysis

All statistical analyses, including descriptive statistics, group comparison test, neutralization curve fitting and visualization, were performed in GraphPad Software v 11.0.0 (Prism). Both parametric and non-parametric analysis were considered, and the appropriate test was selected based on the sample size, data distribution and test assumption. In some analyses, data transformation was conducted to meet the assumption of the statistical tests. Multiple comparison tests were chosen based on the study objective, including whether pairwise comparisons, preselected pairs, or comparisons with a control group were required. The area under the curve (AUC) was calculated for longitudinal data and the corresponding comparison test was performed on the calculated AUC values. Kaplan-Meier survival analysis and log rank test were utilized to analyze survival data. Group means were reported when parametric tests were applied, whereas group medians were reported when nonparametric tests were used.

## RESULTS

### In vitro expression of luciferase or HSV-2 gD using recombinant Ad4 vector

Replication-competent Ad4 expressing luciferase (Ad4-Luc) and HSV-2 gD (Ad4-gD2) were constructed by inserting the respective gene into the E3 region of Ad4 (**Fig. 1A**). Cytopathic effects were observed in A549 cells 3 days after infection with Ad4-Luc (**Fig. 1B,C**) or Ad4-gD2. Lysates from cell cultures infected with Ad4-Luc incubated with luciferin substrate demonstrated a 10^3^-fold increase in luminescence compared to cell lysates from uninfected cells (group median relative luminescence units of 3.0 × 10^7^ versus 1.2 × 10^4^, Kruskal-Wallis test p<0.0001 followed by Dunn’s multiple comparisons test adjusted p = 0.002) (**Fig. 1D**). A band of 55 kDa, corresponding to HSV-2 gD, was detected in lysates from A549 cells infected with Ad4-gD2 by Western blot with anti-gD2 antibody (**Fig. 1E**).

### Kinetics of transgene expression of luciferase in mice after Ad4-Luc inoculation

BALB/c mice were inoculated with Ad4-Luc (1.4 × 10^8^ IFU/ml stock) at various sites at a maximally acceptable volume of inoculum (intranasal: 20 ul per nostril [40 ul total], intramuscular 50 ul per site [100 ul total], gavage: 200 ul and intravaginal 20ul). Mice received 2.7 × 10^6^ IFU of Ad4-Luc in each nostril or Ivag, 6.8 × 10^6^ IFU IM, or 2.2 × 10^7^ IFU OG (**Fig. 2A**). Luminescence was measured using an IVIS Spectrum in vivo Imaging System on live animals on day 3 and 4 as a surrogate for Ad4-Luc early gene expression. The highest luciferase activity was observed on day 4 in the thigh following IM inoculation (group mean of average radiance: 3.0 × 10^4^, range: 2.0 × 10^4^ to 4.4 × 10^4^), followed by luciferase activity in the snout after IN inoculation (group mean of average radiance: 1.0 ×10^4^, range: 3.9 × 10^3^ to 1.8 × 10^4^). Combined day 3 and 4 average radiance values also demonstrated significantly higher luciferase expression in thigh following IM than in snout after IN inoculation (transformed group mean average radiance of 172.8 versus 105.5 respectively (Two-way repeated measures ANOVA on transformed (square-root) average radiance p<0.0001 followed by Šídák’s multiple comparisons test adjusted p=0.0001) (**Fig. 2B**). The group median average radiance after OG or Ivag administration of Ad4-Luc did not exceed the limit of detection (2-fold higher than the group median background average radiance in animals that did not receive Ad4-Luc). A few animals did have values that exceeded the limit of detection, particularly in the snout of those that received Ad4-Luc by OG in bicarbonate with aprotinin, likely due to regurgitation of the virus from the gastrointestinal tract into the nose.

**Fig. 2.**
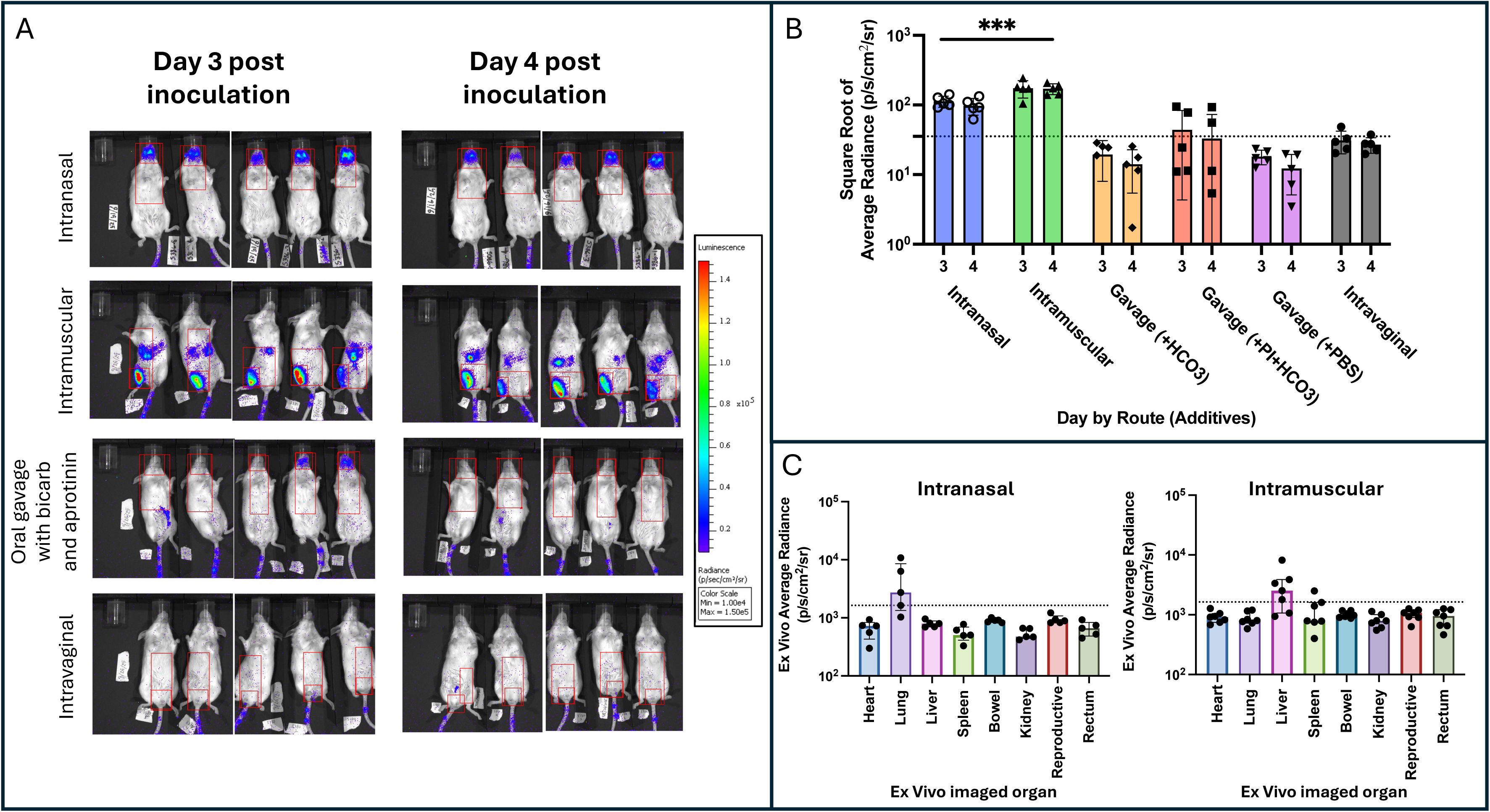
Luciferase activity pattern after different routes of Ad4-Luc inoculation (A) IVIS Spectrum in vivo images of luciferase activity (luminescent signal overlay per color scale) at 3- and 4-days post inoculation of BALB/c mice with Ad4-Luc given via different routes. Intramuscular inoculation was at the right thigh. Each group had 5 mice. Two photos are used to represent 5 mice at each time point for each condition due to limited space in the IVIS imager. (B) Quantitation of luciferase expression 3 and 4 days after Ad4-Luc inoculation. Average radiance from snout after intranasal or oral gavage and from the inoculation site after intramuscular injection or intravaginal inoculation are shown (area encompassed by small red box from IVIS Spectrum in vivo images [see Fig 2A]). Three different diluents were used in the vaccine when administered via the oral gavage route; Phosphate Buffered Saline (PBS), 2% bicarbonate solution (HCO_3_) with or without the protease inhibitor aprotinin (PI). Dashed line represents the limit of detection defined as twice the median background average radiance of an uninfected BALB/c mouse. Imaging sessions were completed for all mice each day but luminescence values below 100 average radiance were not plotted on graph. Bars show mean values with the standard deviation (SD) and individual data points are shown as dots. ***: p=0.0001 (C) Organ-specific ex-vivo luciferase expression four days after intramuscular (right thigh) versus intranasal Ad4-Luc inoculation. Dotted line represents the limit of detection defined as twice the median luminescence value of organs from uninfected mice. Bars show median values with interquartile range and individual data points are shown as dots.

Since luciferase expression was consistently detected above the limit of detection after IN or IM inoculation of Ad4-Luc, luciferase activity was assessed ex vivo in organs removed from mice after IN or IM inoculation. Four days after IN inoculation, luciferase activity was only detected above the limit of detection in the lung (4 of 5 mice) **(Fig. 2C, Supplemental Fig. 2**). Following IM inoculation, luciferase activity was observed above the limit of detection in the liver (5 of 7 mice) (**Fig. 2C; Supplemental Fig. 2**). While expression of luciferase in the lungs after IN inoculation is likely due to direct spread of Ad4-Luc in the respiratory tract, expression in the liver and/or spleen could be due to transport of the virus to these organs by immune cells after IM injection.

Data collected in 19 BALB/c mice from a total of 6 independent experiments were pooled to evaluate the kinetics of luciferase expression after Ad4-Luc IN inoculation over a 3-week period **(Fig. 3A)**. Expression of luciferase in the snout was observed at 7 hours after inoculation. The peak of luciferase expression occurred on day 4, reaching a maximum mean expression level of 1.2 × 10^5^ average radiance and persisted for at least 20 days. Since HAdV is known to replicate better in human cells than mouse cells, we inoculated humanized NCG/NOG mice (that have human hematopoietic cells) IN with Ad4-Luc. The kinetics of luciferase expression in the humanized mice was similar to that in BALB/c mice **(Fig. 3B)**.

**Fig. 3.**
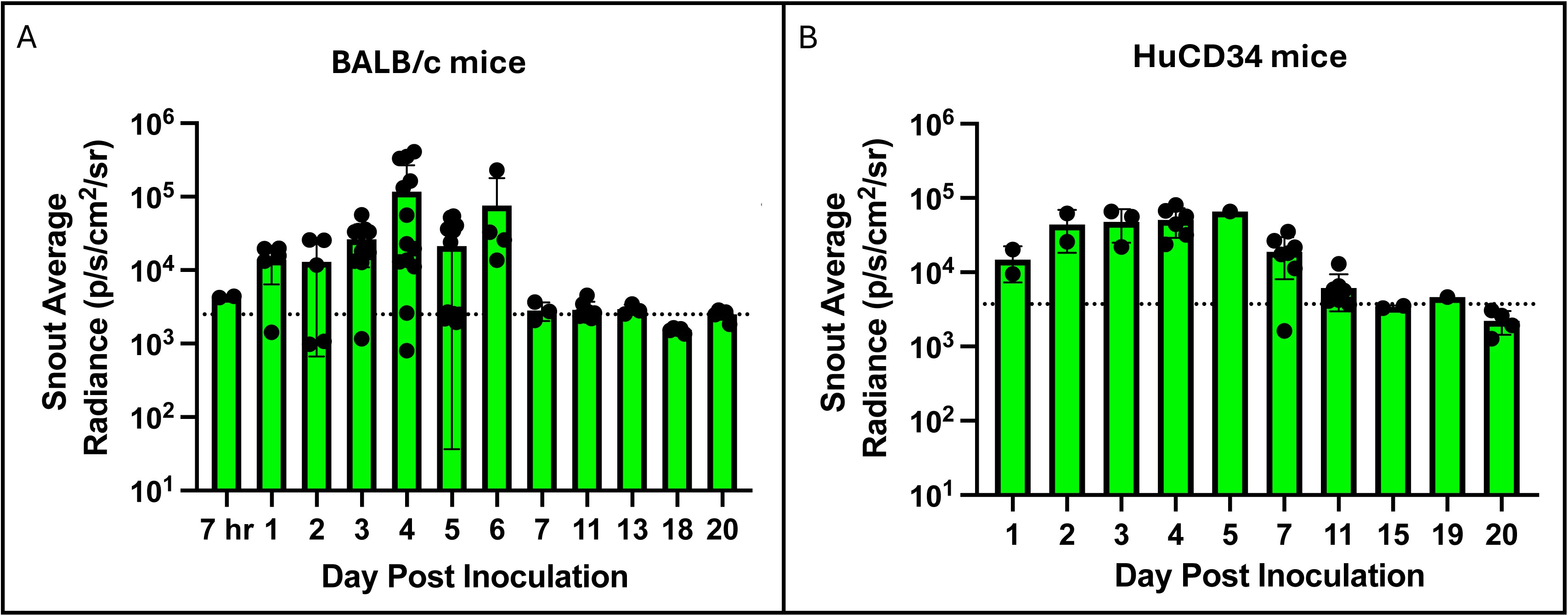
Luciferase snout expression kinetics after intranasal inoculation of BALB/c and humanized mice with Ad4-Luc. (A) Luciferase expression in BALB/c mice from 7 hours to 20 days post inoculation with Ad4-Luc. Each data point represents one mouse imaged during a single imaging session and the figure shows data collected from mice imaged at different times during 6 independent experiments. (B) Luciferase expression in humanized mice (HuCD34) 1 to 20 days after intranasal inoculation with Ad4-Luc. Dotted line represents the limit of detection defined as twice the median background of snout from uninfected mice. Bars show mean values with standard deviations and with individual data points are shown as dots.

### Intranasal inoculation of BALB/c mice, but not humanized mice, with Ad4-Luc reduces luciferase activity upon subsequent Ad4-Luc challenge

The mean luciferase expression in the snout of BALB/c mice inoculated with Ad4-Luc decreased an average of 23-fold between the first and second intranasal inoculation (mean average radiance 8.1 × 10^4^ vs. 3.5 × 10^3^ three days after first and after the second inoculation, respectively; paired t-test p = 0.005) (**Fig. 4A,B**). In a parallel experiment, we observed an increase in mean Ad4-Luc ID_50_ neutralizing serum titers (dilution of serum antibody needed to inhibit viral entry by 50%) from18.6 on day -2 to 126.0 on day 94; paired t-test p=0.032) of five BALB/c mice that received two intranasal doses of Ad4-Luc (**Supplemental Fig. 3).** In contrast to inoculation of BALB/c mice, robust luciferase activity was observed after repeat intranasal inoculation of humanized mice with Ad4-Luc. The mean average snout radiance remained similar three days after the second intranasal Ad4-Luc dose compared to three to four days after the initial dose (mean average radiance 5.0 × 10^4^ vs. 4.9 × 10^4^, respectively; paired t-test p= 0.957) (**Fig. 4A,B**). Since humanized mice are immunocompromised, they likely have an absent or impaired immune response to the Ad4 vector and therefore reinfection with Ad4 achieves a similar level of infection as the primary infection. Taken together, these experiments suggest that an immune response to the Ad4 vector reduces infection and/or replication after the second dose of the vector in immunocompetent BALB/c mice.

**Fig. 4.**
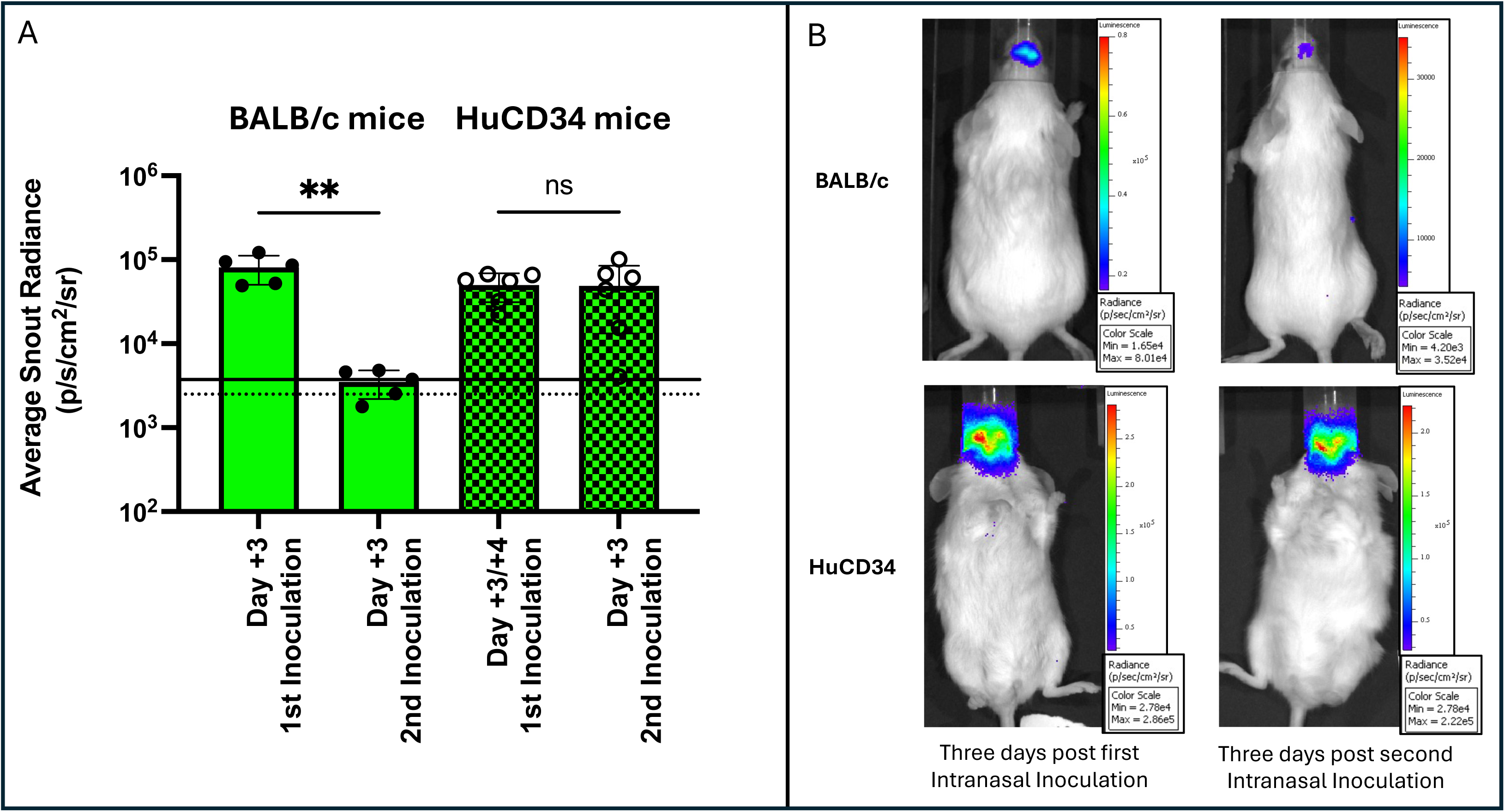
Luciferase expression decreases after multiple intranasal doses of Ad4-Luc in BALB/c mice, but not in humanized mice. (A) Solid line and dotted line represent the limit of detection of average snout radiance for HuCD34 mice and for BALB/c respectively. Bars show mean values with standard deviations and individual data points are shown as dots. Five BALB/c and six HuCD34 mice were used in the experiment. ** p=0.005; ns: not significant. (B) Sequential photographs of a BALB/c and humanized mouse 3 days after the first and 3 days after the second intranasal inoculation of Ad4-Luc. Differences in luciferase expression can be observed as snout luminescence (luminescent signal overlay per color scale).

### Cidofovir inhibits replication of Ad4-Luc in mouse lung tissue

To determine whether Ad4-Luc replicates in mice after IN inoculation, luciferase expression and quantification of adenovirus DNA in the lungs were evaluated in animals that received intraperitoneal cidofovir daily for 4 days beginning on the day of Ad4-Luc infection. In vivo imaging of snout luciferase activity was not different between cidofovir treated and untreated mice on daily imaging between day 1 and day 4 post inoculation with Ad4-Luc (mean area under the curve (AUC) of 4.7 × 10^4^ versus 3.8 × 10^4^ respectively, paired t-test p=0.321) **(Fig. 5A).** Luciferase activity in ex vivo lung tissue extracted on day 4 after intranasal inoculation with Ad4-Luc was not statistically different in mice that had received cidofovir compared to cidofovir naïve mice (mean average radiance per well containing whole right lung of 2092 versus 2088, respectively; Welch’s t-test p= 0.995) **(Fig. 5B)**. In contrast, adenovirus DNA was decreased in lung tissue of cidofovir treated mice vs. untreated mice (mean adenovirus DNA copies per mg of lung tissue of 855.6 versus 2581, respectively; Welch’s t–test p=0.033) **(Fig. 5C).** These data suggest that while cidofovir did not limit infection, it did diminish adenoviral DNA synthesis in vivo.

**Fig. 5.**
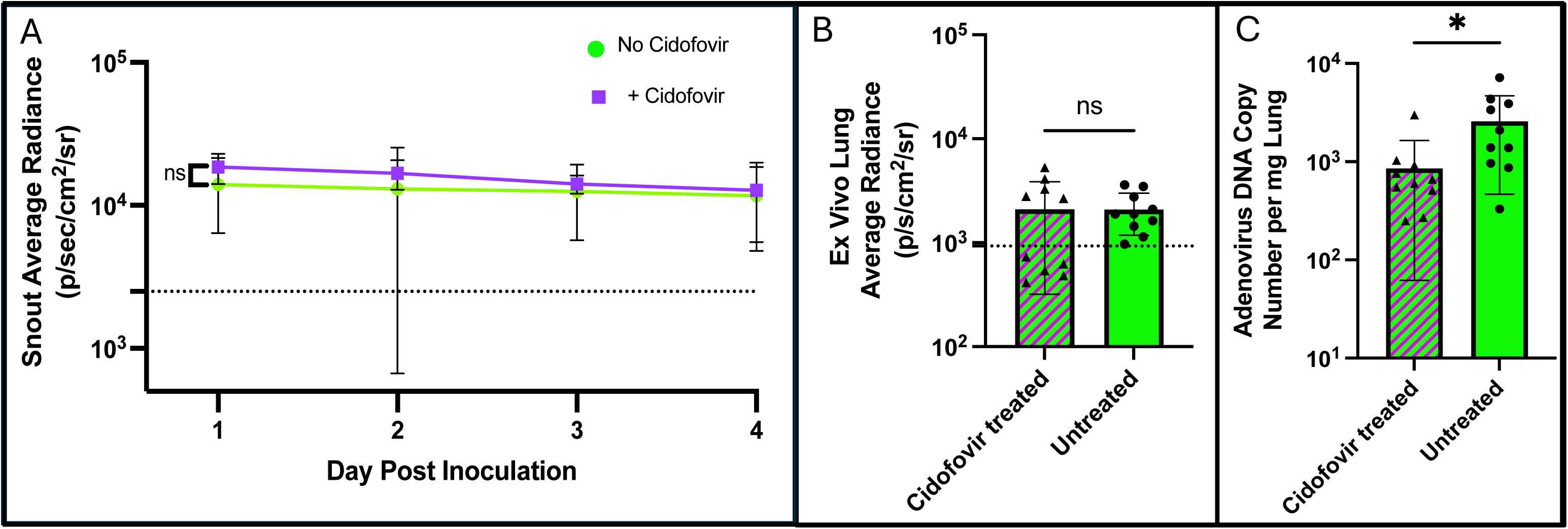
Cidofovir anti-viral therapy decreases adenovirus DNA in lung tissue 4 days after inoculation. (A) In vivo snout luminescence was calculated daily from day 1 to day 4 after Ad4-Luc intranasal infection in cidofovir treated (purple striped) and untreated mice (solid green) mice. Error bars represent the mean values with standard deviations. Five male mice were used in each group. ns: not significant. (B) Luciferase expression of ex vivo lung tissue (whole right lung en bloc) in cidofovir treated (purple striped) and untreated (solid green) mice was assessed via luminescence (average radiance of well containing lung) in the IVIS system 4 days after intranasal inoculation with Ad4-Luc. Bars show mean values with standard deviations and individual data points are shown as dots. ns: not significant. (C) Quantitative PCR for adenovirus DNA in the left lung after necropsy. All samples were run on the same day with samples for the standard curve. Bars show mean values with standard deviations and individual data points are shown as dots. Twenty mice (10 male, 10 female) from two independent experiments were used in panels B and C. *: p=0.033.

### Mice vaccinated with Ad4-gD2 intranasally develop higher titer anti-HSV-2 gD antibody responses than mice vaccinated with Ad4-gD2 by other routes

HSV-2 causes genital, oral, and ocular mucocutaneous infections in humans and no vaccines have been licensed to prevent HSV-2 disease. A mouse model of genital HSV-2 has been extensively used for testing the efficacy of HSV-2 vaccines ^36,37^. To test the Ad4 replication-competent vector as a vaccine against HSV-2 infection in mice, we inoculated BALB/c mice with Ad4-gD2. HSV-2 gD (gD2) was chosen as the immunogen because it is a major target of HSV-2 neutralizing antibodies in humans and the viral glycoprotein that binds to the host cellular receptors (nectin-1 and HVEM)^36^. Mice were vaccinated twice, 3 weeks apart with Ad4-gD2 by the IN (5.0 × 10^11^ IFU per dose), OG (1.8 × 10^12^ IFU per dose), or Ivag (2.0 × 10^11^ IFU per dose) route or with soluble gD2 protein (5 ug per dose) mixed with Sigma Adjuvant Systems (SAS) by the IM route. One group of mice received two IM injections of phosphate buffered saline (PBS) with SAS and served as a control. Serum was obtained for HSV-2 gD ELISA and virus neutralizing titers 7 days after the second vaccination. Serum anti-HSV-2 gD IgG endpoint titers were highest in mice vaccinated IM with soluble gD (group mean: 4.2 × 10^5^), followed by Ad4-gD2 administered IN (group mean: 1.1 × 10^5^), OG (group mean: 7.6 × 10^3^) and then Ad4-gD2 administered Ivag (group mean: 2.4 × 10^3^) (**Fig. 6A**). The geometric mean titer of each Ad4-gD group was significantly lower than the soluble gD IM group: Ad4-gD2 IN (adjusted p = 0.005), OG (adjusted p = 0.0003) and Ivag (adjusted p < 0.0001) (lognormal Brown-Forsythe and Welch ANOVA test p < 0.0001 followed by Holm-Šídák’s multiple comparisons test). However, serum HSV-2 neutralizing antibody titers were similar in mice that received Ad4-gD2 IN and mice that received soluble gD IM (group mean: IC_50_ 71.8 versus 79.4; Welch’s t test p = 0.494) (**Fig. 6B**).

**Fig. 6.**
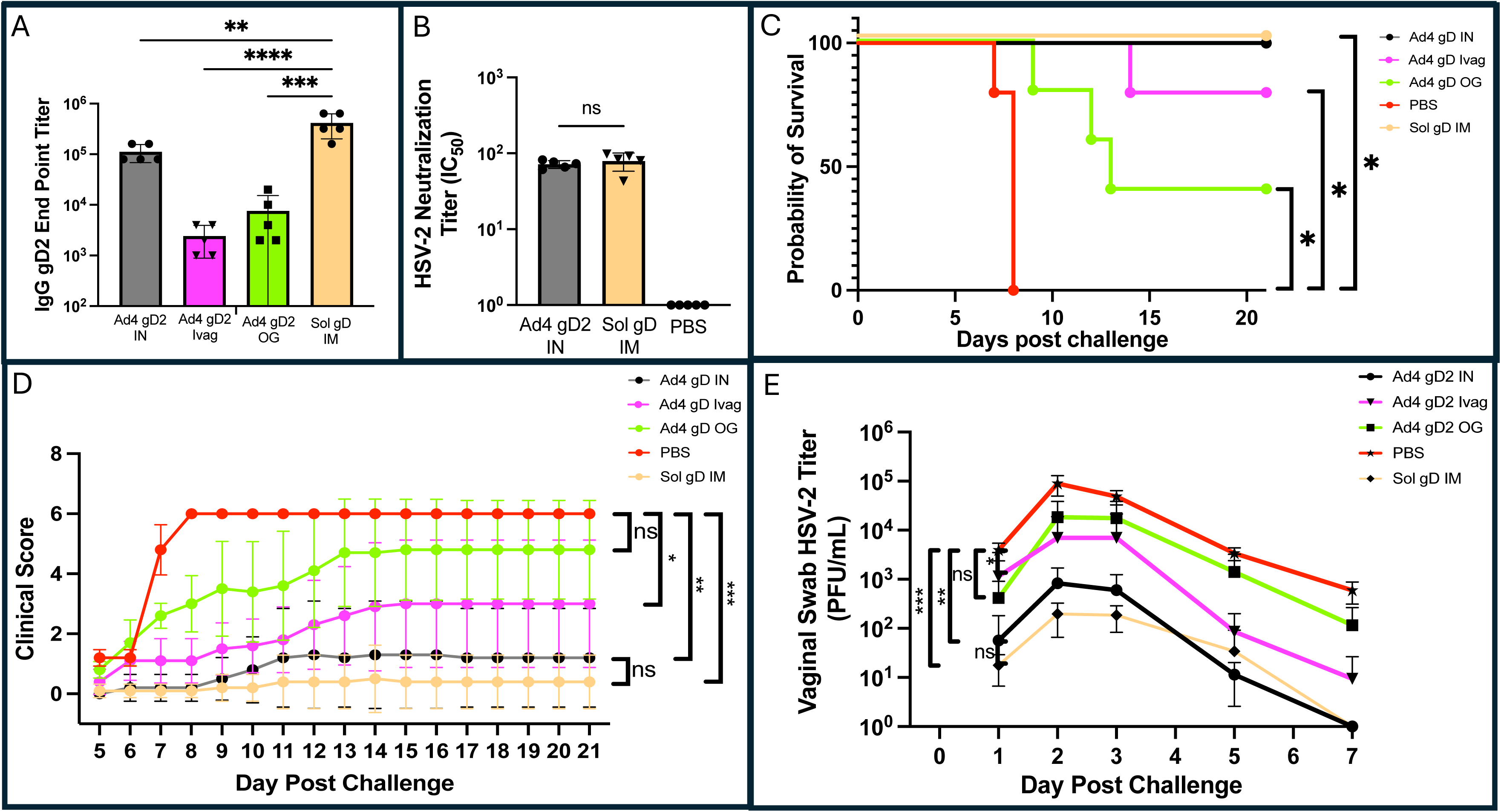
Intranasal Ad4-gD2 inoculation elicits HSV-2 antibodies and protects mice from disease and decreases HSV-2 shedding after intravaginal challenge with HSV-2. (A) HSV-2 gD2 ELISA titers of sera from mice after two doses of either Ad4-gD2 (given IN, Ivag, or by oral gavage) or soluble gD2 (given IM). Bars show mean values with standard deviations with individual data points shown as dots. **: p=0.005; ***: p=0.0003; ****: p < 0.0001. (B) Serum HSV-2 neutralization titers in mice that received two doses of either Ad4-Luc (IN) or soluble gD (IM); PBS treated mice are a negative control. Sera was obtained 1 week after the second dose of vaccine. Bars show mean values with standard deviations with individual data points shown as dots. ns: not significant. (C) Survival curves of mice inoculated with Ad4-gD2 (given by various routes), soluble gD (IM), or PBS after intravaginal challenge with 6.4 × 10^4^ PFU of HSV-2 given intravaginally. Kaplan-Meier survival analysis was conducted to assess survival of mice after vaginal HSV2 challenge among 5 groups. *: p = 0.014._(D) Group disease scores of mice after intravaginal challenge with HSV-2. Mean daily scores with standard deviations for each group at each day post-challenge. Each vaccine group had 5 mice. *: p=0.028; **: p=0.005; ***: p=0.0003; ns: not significant. (E) Vaginal shedding of HSV-2 in mice inoculated with Ad4-gD2 (given by various routes), soluble gD (IM), or PBS after intravaginal HSV-2 challenge. Group mean values are shown with standard deviations. HSV-2 was titrated on Vero cells. Bars show group means of HSV-2 titer with standard deviations. Each vaccine group consisted of 5 mice.*: p = 0.048; **: p = 0.003; ***: p = 0.0002; ns: not significant.

### Mice vaccinated with Ad4-gD2 intranasally have increased survival after HSV-2 intravaginal challenge and decreased HSV-2 shedding compared with animals vaccinated by other routes

Ad4-gD2 vaccinated mice were challenged Ivag with HSV-2 14 days after receiving the second dose of vaccine. Mice vaccinated IN with Ad4-gD2 or IM with soluble gD2 had 100% survival after challenge, compared to 0% survival in the PBS group. In contrast, 80% of mice inoculated Ivag with Ad4-gD2 and 40% vaccinated OG with Ad4 survived after challenge (**Fig. 6C**). The survival curves of each vaccine groups were significantly different from the PBS group (log-rank Mantel-Cox test p<0.0001 followed by Holm-Šídák’s multiple comparisons test, all adjusted p = 0.014). Disease scores generally reflected the trend in survival after challenge; mice vaccinated with Ad4-gD IN had a group mean disease score of 0.9 (range: 0 to 3.5) on day 21 post challenge and mice vaccinated with soluble gD IM had a disease score of 0.3 (range: 0 to 2.5) (**Fig. 6D**). In contrast, mice vaccinated with Ad4-gD Ivag had a day 21 group mean disease score of 2.2 (range: 0 to 6), and mice receiving Ad4-gD OG had a day 21 group mean disease score of 3.9 (range: 0.5 to 6). The AUC of disease scores were significantly different among vaccine groups (Brown-Forsythe and Welch ANOVA tests p < 0.0001). Specifically, the mean AUC of disease scores from three of the vaccine groups were significantly lower than PBS group: Ad4 gD IN (adjusted p = 0.005), Ad4 gD Ivag (adjusted p = 0.028) and soluble gD IM (adjusted p = 0.0003). However, the AUC of disease scores between Ad4 gD OG and PBS (mean AUC: 62.9 versus 87.6; adjusted p = 0.223) and between Ad4 gD IN and soluble gD IM (mean AUC: 14.9 versus 5.1; adjusted p = 0.864) were not significantly different (Dunnett’s T3 multiple comparisons test).

HSV-2 shedding was measured as an indicator of the ability of Ad4-gD2 vaccine to block HSV-2 infection and virus replication. Viral shedding after HSV-2 challenge was observed in all mice, peaking on day 2 post challenge (**Fig. 6E**). Vaginal shedding was lowest in mice vaccinated with soluble gD IM (day 2 group mean titer: 2.0 × 10^2^ PFU) and slightly higher in Ad4-gD2 IN vaccinated animals (day 2 group mean titer: 8.3 × 10^2^ PFU). In contrast, mice vaccinated with Ad4-gD2 by the Ivag route had higher levels of HSV-2 shedding (day 2 group mean titer: 7.0 × 10^3^ PFU), followed by mice vaccinated with Ad4-gD2 by the OG route (day 2 group mean titer 1.9 × 10^4^ PFU), followed by the PBS group (day 2 group mean titer 9.0 × 10^4^ PFU). Differences between AUC of HSV-2 shedding over 7 days among vaccine groups were assessed using lognormal Brown-Forsythe test followed by Holm-Sidak’s multiple comparisons test. The differences in virus shedding between mice that were given PBS and animal vaccinated with soluble gD (adjusted p=0.0002), Ad4-gD2 IN (adjusted p=0.003), and Ad4-gD2 Ivag (adjusted p=0.048) were all significant. Mice immunized with IN Ad4-gD2 demonstrated a similar level of HSV-2 shedding over the 7-day period post challenge compared to soluble gD (geometric mean HSV-2 shedding titer AUC: 1017 versus 423, respectively; adjusted p = 0.277). Immunization with Ad4-gD2 IN reduced HSV-2 shedding 108-fold, compared to PBS, on day 2 following Ivag HSV-2 challenge. Thus, IN Ad-gD2 was sufficiently immunogenic in mice to induce serum neutralizing antibody responses, prevent mortality, reduce disease scores, and inhibit shedding after Ivag challenge with HSV-2

## DISCUSSION

In this paper we evaluated Ad4 transgene expression kinetics in mice and the use of the virus as a vaccine vector to protect mice from HSV infection and disease. Ad4 has been used as part of an Ad4/Ad7 bivalent vaccine in the U.S. military for decades^10^. Furthermore, adenovirus vectors have been shown to generate potent immune responses against a myriad of other pathogens including influenza virus, coronavirus, hepatitis B virus, hepatitis C virus, Ebola virus, and HIV^11,31,38–42^. Human adenoviruses are generally thought to exhibit species-specific replication, with minimal replication in animals, including non-human primates. Cotton rats and Syrian hamsters show semi-permissive replication of group C human adenoviruses and have historically been used as model systems studying anti-adenovirus therapeutics^43,44^. While there is generally limited adenovirus replication in most mouse tissues and cell lines, there are reports of intravenously and intranasally administered Ad5 inducing disease in BALB/c or CBA and C57BL/6N mice respectively^28,45 46^ . The abundance of immunologic reagents, and ease of working with these animals, make mice the favored model system for studying immune responses in preclinical development of vaccines.

We evaluated various sites of Ad4 inoculation, including mucosal surfaces, since there is increasing evidence that development of mucosal immunity is important for both short- and long-term protection from pathogens that infect the mucosa. While intramuscular vaccines effectively induce systemic pathogen-specific immune responses, mucosal vaccines induce localized antibody and cellular immune responses^47^. IgG appears to play an important role to protect the reproductive tract mucosa, specifically vaginal tissue from HSV-2 infection^48^. Both intramuscular and oral polio vaccination prevent poliomyelitis, but the oral polio vaccine induces localized immune responses in the gastrointestinal tract and more effectively reduces shedding, to limit person-to-person transmission^49^. In addition, induction of anti-hemagglutinin and neuraminidase antibodies in the nasal mucosa from intranasal live influenza vaccination (FluMist^®^) recapitulates localized respiratory immune response to natural influenza infection^50^. Nasopharynx-associated lymphoid tissue (NALT) in rodents, analogous to Waldeyer’s ring lymphoid tissue in humans, contains lymphoid cells that direct IgA+ B-cell class switching as well as stimulate the expansion of local cytotoxic T cells in response to viral infection. In response to intranasal vaccination, NALT tissue appears to induce immune responses in both the respiratory and reproductive tract^51^. We investigated three mucosal routes- nasal, oral and vaginal- for Ad4 administration in the mouse model. We found that both intranasal and intramuscular administration of Ad4-Luc resulted in prolonged expression of luciferase at the site of inoculation, while intravaginal and orogastric administration were not effective to express luciferase. It is not surprising that orogastric Ad-Luc was ineffective as the stomach is acidic and oral administration of the military Ad4/Ad7 bivalent vaccine requires an enteric coating which was not used in our experiments^7^. Cells in the vaginal tract lack apical expression of the coxsackie and adenovirus receptor (CAR)^52^ and while some adenovirus serotypes infect the genital tract, Ad4 is not a frequent cause of genital infection.^53,54^

Different routes of delivery of adenovirus vectors result in varying degrees of infection of local tissues and sometimes dose-limiting organ toxicity^55–58^. We found differences in luciferase expression in different organs of mice inoculated with Ad4-Luc by the intranasal route and by the intramuscular route. Intranasal inoculation resulted primarily in luciferase activity in the upper respiratory tract and the lungs, while intramuscular inoculation resulted in luciferase activity in the muscle as well as the liver, indicating hematogenous spread of the virus. Due to safety concerns associated with dissemination of adenovirus to various organs and the history of liver toxicity in non-human primate models and patients with systemically administered adenoviral vectors,^57,58^ we focused on intranasal Ad4 administration for most of our subsequent experiments.

We evaluated the duration of transgene expression in mice that received intranasal Ad4-Luc. We found that both immunocompetent BALB/c mice and immunocompromised humanized mice had luciferase activity for approximately 20 days in the snout. There was less individual variability in humanized mice than BALB/c mice. In addition, BALB/c mice showed lower peak snout luciferase activity after a second dose of Ad4-Luc, while humanized mice had robust peak snout luciferase activity with a second dose similar to that seen with the initial inoculation. This suggests an immune response may be limiting Ad4 replication after a primary or secondary infection of immunocompetent mice. Taken together, our results indicate that Ad4 vectors can achieve transgene expression for more than 1 week.

To evaluate whether Ad4 replicates in mice as it does in humans, and therefore whether the mouse model might be predictive for preclinical studies of vaccines, we performed mouse infections in the presence or absence of cidofovir. Cidofovir inhibits adenovirus replication by becoming incorporated into viral DNA resulting in DNA chain termination and by direct inhibition of the viral polymerase^59^. Cidofovir has been shown to inhibit replication of mouse adenovirus in mice^60^ and human adenovirus in hamsters^61,62^. Cidofovir treatment did not significantly change luciferase activity when BALB/c mice were imaged, suggesting the majority of transgene expression in mouse snouts came from the initial Ad4-Luc inoculum and not progeny virus. In addition to limited nasal replication, this result may also be a consequence of poor nasal tissue penetrance of cidofovir after intraperitoneal dosing. A study in rabbits showed that the cidofovir accumulated in the lungs at levels that were 3-fold higher than in the skin at 6 and 24 hours following intravenous administration^63^. Consistent with our findings for live mouse imaging, we found similar lung luciferase activity observed in lungs removed from cidofovir treated mice compared to untreated controls. In contrast, we found that cidofovir significantly reduced adenovirus replication in the lungs as evidenced by reduced Ad4 DNA in the lungs of cidofovir treated mice compared with untreated controls. This implies that there is replication of Ad4 in the lungs of healthy BALB/c mice after intranasal infection. These results emphasize the potential benefit of replication-competent adenovirus compared with replication-incompetent virus in that the former can persist and induce a more persistent immune response. Stimulation of host immune responses with replication-competent Ad4 expressing influenza H5 hemagglutinin produced potent, long-lasting adaptive immune responses to H5 hemagglutinin in humans^13^ .

Finally, we tested whether the immune response generated from a replication-competent Ad4 vector with an inserted transgene in mice via different routes of mucosal vaccination could provide significant protection from challenge with HSV-2. We inoculated mice twice with Ad4 containing HSV-2 gD (Ad4-gD2) via the intranasal, oral, or intravaginal route and compared immune responses and protection from disease with mice inoculated with soluble HSV-2 gD with adjuvant given IM, an extensively studied HSV-2 vaccine candidate in human. Intranasal, but not orogastric or intravaginal inoculation with Ad4-gD2 elicited nearly similar HSV-2 gD antibody titers as soluble HSV-2 gD given IM. Neutralizing antibody titers against HSV-2 generated by intranasal Ad4-gD2 were equivalent to soluble HSV-2 gD IM. Furthermore, intranasal, but not orogastric or intravaginal inoculation with Ad4-gD2, resulted in 100% survival, reduced disease scores, and reduced HSV-2 vaginal shedding to a similar degree as soluble HSV-2 gD IM after challenge with HSV-2. Other studies have produced similar results using a two- or three-dose IM regimen of replication-incompetent Ad35 or Ad5 respectively expressing HSV-2 gD^64,65^. Notably, both groups observed additional HSV-2 protection when an additional immunogen (HSV-2 gB or UL25) was added to gD as either an additional recombinant adenovirus or a fusion protein respectively. Other investigators have found that increasing gD antigen expression by using a recombinant Ad5 with four copies of HSV-1 gD increased immunogenicity versus the same vector with a single copy of HSV-1 gD^66^. Considering the permissibility of Ad4 in human hosts, we would expect Ad4-gD2 replication, transgene expression, and immunogenicity to be higher in humans than in mice. Prior exposure to wild-type Ad5 infection in cotton rats appears to limit their response to a recombinant Ad5 vector expressing bovine herpesvirus gD^67^. Similarly, preexisting Ad4 immunity in humans would likely reduce recombinant Ad4 replication and subsequent immunogenicity. While we evaluated humoral immune responses to Ad4-gD2, other investigators observed localized CD4+ and CD8+ T cell responses that correlated with reduced recurrent genital lesions and decreased viral shedding in guinea pigs latently infected with HSV-2 that were subsequently vaccinated with a replication incompetent Ad5 expressing gD2^68^. In our study, intravaginal vaccination with Ad4-gD2 likely induced localized gD-specific T-cell responses that contributed to the protection from lethal HSV-2 intravaginal challenge, despite low anti-gD antibody titers in the blood.

Our study has several limitations. First, replication of Ad4 was limited in mice. We used a higher titer inoculum of Ad4 expressing HSV-2 gD2 in mice (10^11^ IFU) compared to the dose of Ad4 expressing influenza H5 hemagglutinin in humans (10^8^ IFU)^13^ to overcome the limited replication of Ad4 in mice. While safety profiles will need to be established for individual recombinant Ad4 candidates, we can expect more robust immune responses in humans than mice, likely requiring lower doses to receive the same or better effect. Second, the potential use of replication-competent Ad4 vectors raises the concern for the development of adenovirus disease after vaccination, especially in immunocompromised hosts^69^. While any potential Ad4-based vaccine will need to undergo rigorous safety testing before being adopted for clinical use, the long track record of safety with the bivalent Ad4/Ad7 vaccine in the military provides some reassurance. In addition, clinical studies of a recombinant Ad4 expressing influenza HA showed that it was well tolerated^12,13^. Third, the mouse model does not recapitulate many of the features of human HSV-2 disease; HSV-2 is typically lethal in mice and in mice that survive infection, the virus does not reactivate like it does in humans.

In summary, we found that Ad4 vectors administered at different sites of inoculation resulted in different levels of transgene expression in various tissues of BALB/c mice. These results may be helpful in designing Ad4 vector systems to generate tissue-specific immunity while limiting toxicity. Furthermore, we found that intranasal administration of replication-competent Ad4 (Ad4-Luc) to mice resulted in modest replication of the virus in the respiratory tract. This implies that mice can be used to initially evaluate Ad4 vectored vaccines. Finally, we demonstrated that an Ad4 vector expressing HSV-2 gD induced a robust anti-HSV-2 immune response that rescued mice from a lethal HSV-2 challenge. Taken together, these data highlight the usefulness of studying Ad4-based vectors in the BALB/c mouse model and demonstrate the potential for using replication-competent Ad4 vectors for future vaccines.

## Funding

This research was supported in part by the Intramural Research Program of the National Institutes of Health (NIH). The contributions of the NIH author(s) were made as part of their official duties as NIH federal employees, are in compliance with agency policy requirements, and are considered Works of the United States Government. However, the findings and conclusions presented in this paper are those of the authors and do not necessarily reflect the views of the NIH or the U.S. Department of Health and Human Services, nor does mention of trade names, commercial products, or organizations imply endorsement by the U.S. Government. This project has been funded in part with Federal funds from the National Institute of Allergy and Infectious Diseases (NIAID), National Institutes of Health, Department of Health and Human Services under BCBB Support Services Contract HHSN316201300006W/75N93022F00001 to Guidehouse Digital.

## Disclosure

The authors of this manuscript have no conflicts of interest to disclose.

## Acknowledgements

We would like to thank the excellent veterinary and technical support from NIAID’s Comparative Medicine Branch specifically Tierra Hunt, Olivia Tuminelli and the staff of the 14BS animal facility. We would also like to thank Alexander Stewart of the Visual and Medical Arts at the Research Technologies Branch of NIAID for assistance with the Ad4 virus particle and genome diagram of Figure 1A.

## SUPPLEMENTAL MATERIAL

**Supplemental Fig. 1.**
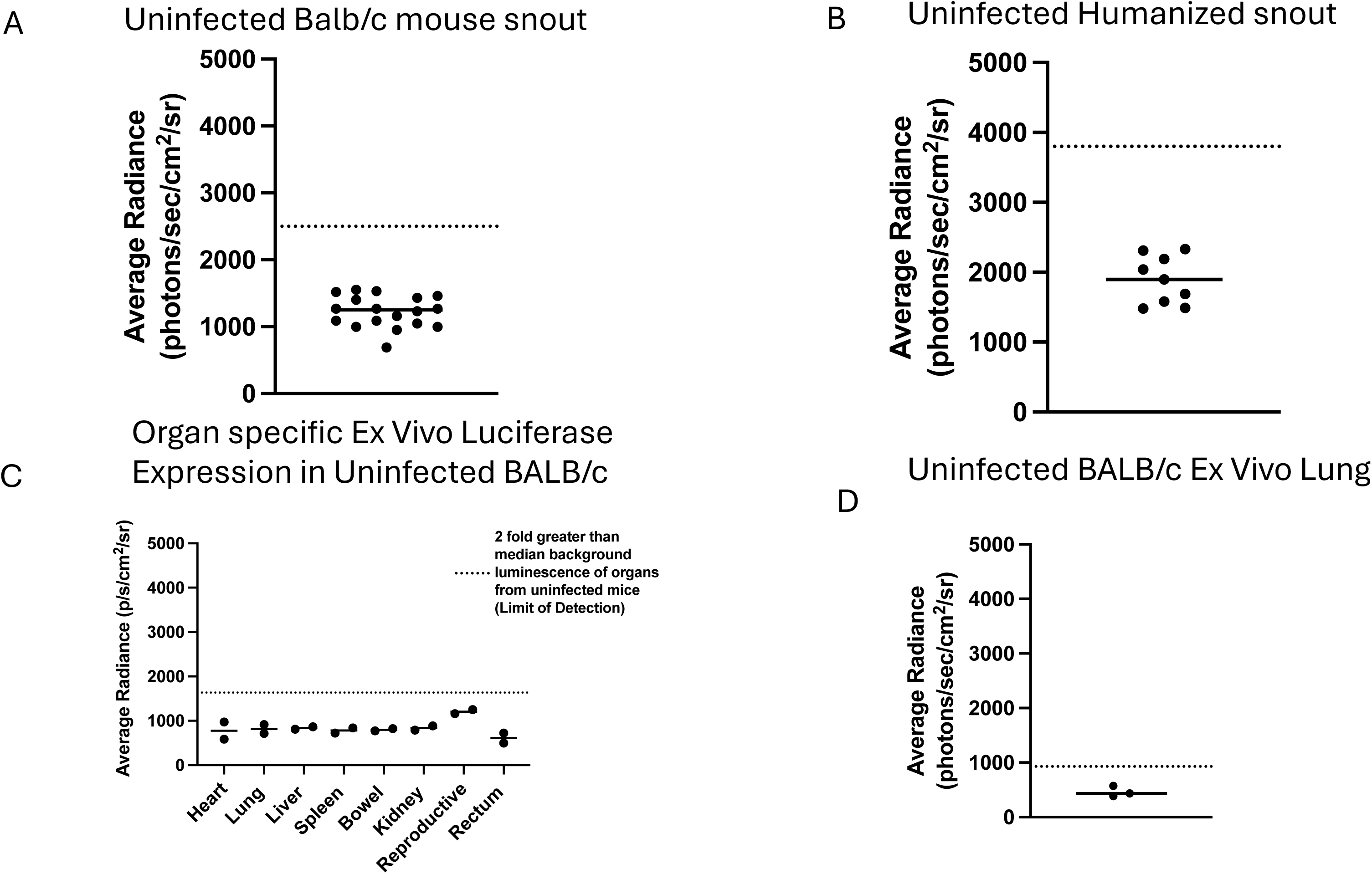
Limit of detection calculation for selected IVIS Assay (A) Average radiance of the whole body of three uninfected BALB/c mice (1 female, 2 male) imaged either supine (n=2) or prone (n=1). Each point represents an independent imaging session. Median average radiance value (1,250 average radiance) was used to determine the limit of detection of the assay. (B) Average radiance of the whole body of two uninfected HuCD34 mice (2 female) imaged either supine (n=1) or prone (n=1). Each point represents an independent imaging session. Median average radiance value (1,900 average radiance) used to determine the limit of detection of the assay. (C) Organ-specific ex vivo background bioluminescence from uninfected mice (n=2). Median average radiance (818 average radiance) (D) Median average radiance from uninfected ex vivo lungs (434 average radiance) from uninfected mice (n=3). Dotted lines in all panels represent the limit of detection for each assay defined as twice the median average limit of detection.

**Supplemental Fig. 2.**
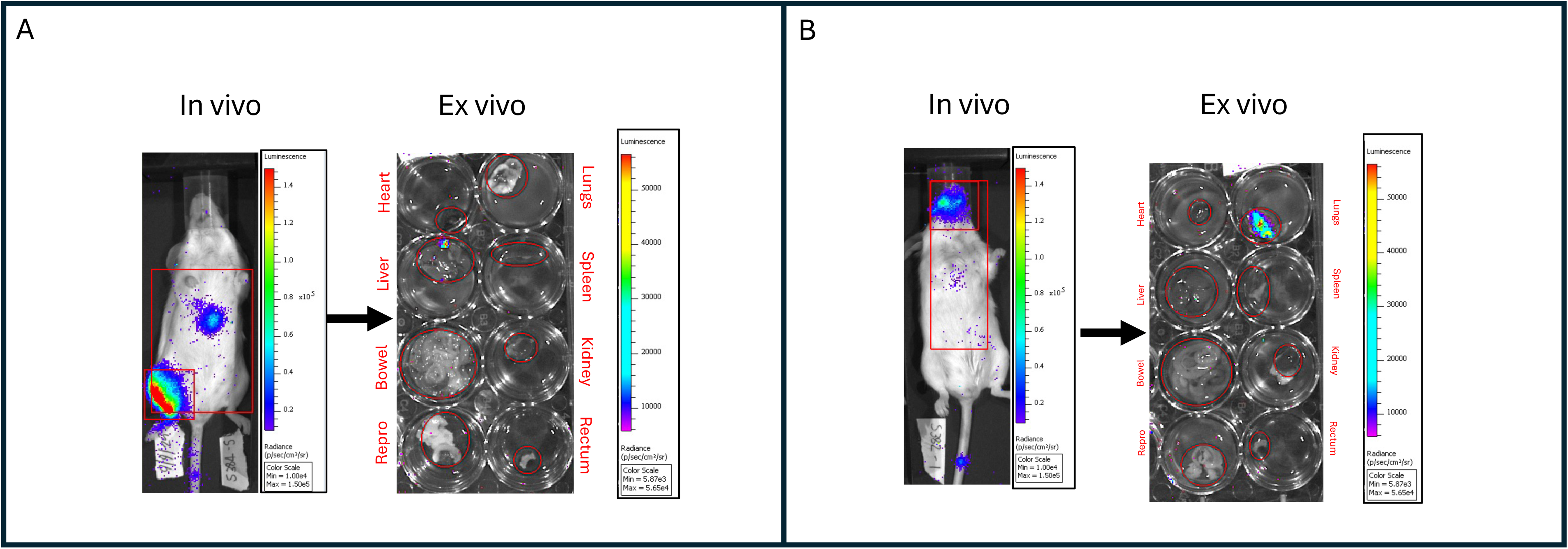
Example of in vivo and ex vivo IVIS imaging of BALB/c mice 4 days after inoculation with Ad4-Luc. (A) In vivo versus ex vivo comparison of organ specific luciferase expression 4 days after intramuscular inoculation with Ad4-Luc (6.8 ×10^6^ total IFU in the right thigh). Luciferase signal is detected in the liver. (B) In vivo versus ex vivo comparison of organ specific luciferase expression four days after intranasal inoculation with Ad4-Luc (5.4 ×10^6^ total IFU). Luciferase signal is detected in the lungs.

**Supplemental Fig. 3.**
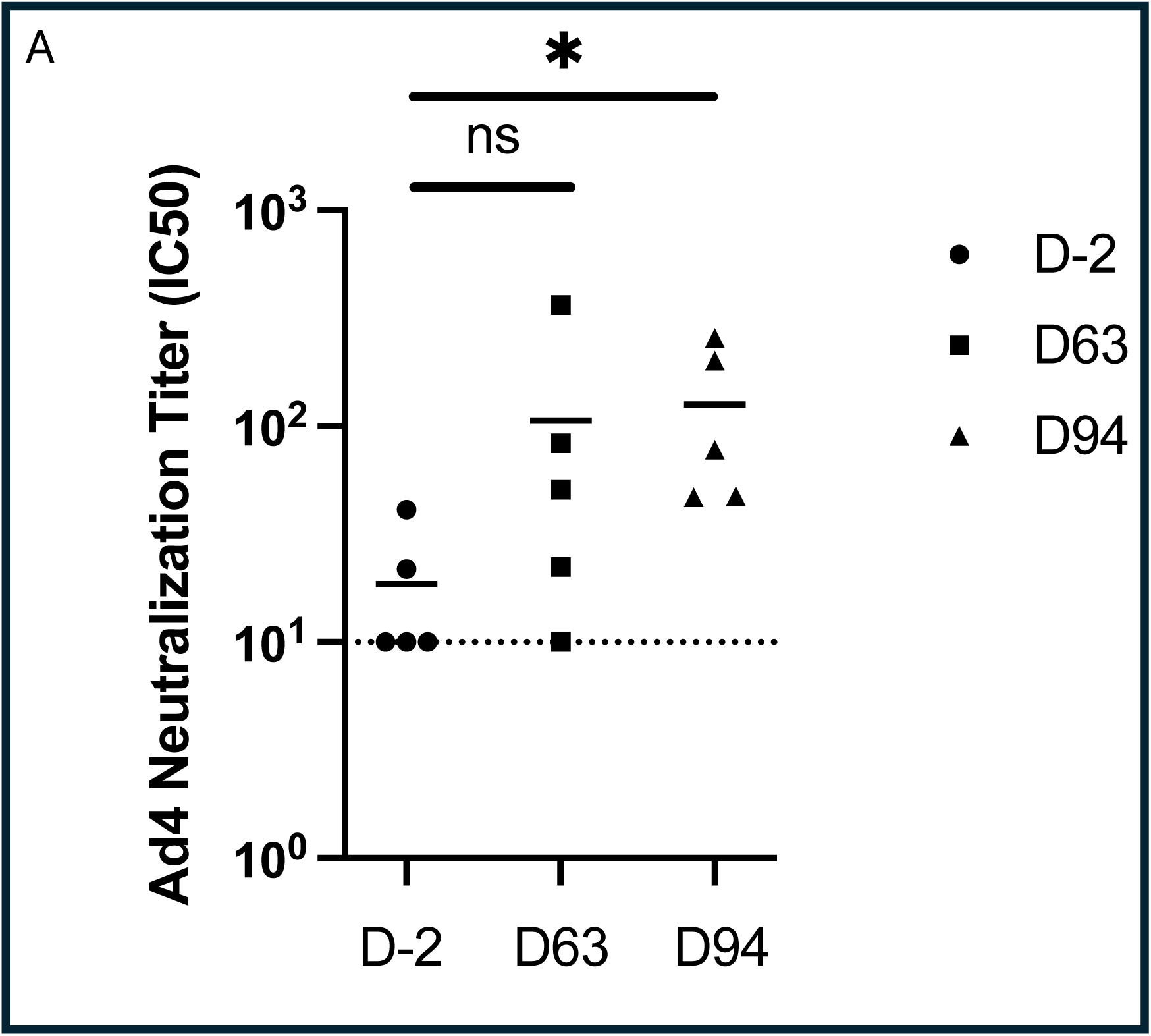
Ad4 neutralizing antibody titers in the serum after two intranasal doses Ad4-Luc. Ad4 serum neutralization titers (50% inhibitory concentration, IC_50_) of mice immunized with two intranasal doses of Ad4-Luc (given on day 0 and day 42) at days -2, 63 and 94. Horizontal lines indicating the group means. Dashed line is limit of detection of assay. *: p = 0.032; ns: not significant.

